# Cilia loss on distinct neuron populations differentially alters cocaine-induced locomotion and reward

**DOI:** 10.1101/2023.06.22.546096

**Authors:** Thomas Everett, Tyler W. Ten Eyck, Chang-Hung Wu, Amanda L. Shelowitz, Sofia M. Stansbury, Ally Firek, Barry Setlow, Jeremy C. McIntyre

## Abstract

Neuronal primary cilia are being recognized for their role in mediating signaling associated with a variety of neurobehaviors, including responses to drugs of abuse. Primary cilia are microtubule-based organelles that project from the surface of nearly all mammalian cells, including neurons. They function as signaling hubs and are enriched with a diverse array of GPCRs, including several known to be associated with motivation and drug-related behaviors; however, our understanding of how cilia regulate neuronal function and behavior is still limited. The objective of the current study was to investigate the contributions of primary cilia on specific neuronal populations to behavioral responses to cocaine. To test the consequences of cilia loss on cocaine-induced locomotion and reward-related behavior, we selectively ablated cilia from dopaminergic or GAD2-GABAergic neurons in male and female mice. Cilia ablation on either population of neurons failed to significantly alter acute locomotor responses to cocaine at a range of doses. With repeated administration, mice lacking cilia on GAD2-GABAergic neurons exhibited greater locomotor sensitization to cocaine compared to wild-type littermates, whereas mice lacking cilia on dopaminergic neurons exhibited reduced locomotor sensitization to cocaine at 10 & 30mg/kg. Mice lacking cilia on GAD2-GABAergic neurons showed no difference in cocaine conditioned place preference (CPP), whereas mice lacking cilia on dopaminergic neurons exhibited reduced CPP compared to wild-type littermates. Combined with previous findings using amphetamine, our results show that behavioral effects of cilia ablation are cell-and drug type-specific, and that neuronal cilia contribute to modulation of both the locomotor-inducing and rewarding properties of cocaine.

## Introduction

Despite advancements in our understanding of the neural systems that mediate drug use, rates of substance use disorders continue to increase rapidly in the United States, and we lack robust and efficacious targets in the brain for pharmacotherapy of substance use disorders (Kwako *et al*. 2016; Potenza *et al*. 2011; Volkow and Li 2005). Recently, neuronal primary cilia have emerged as significant regulators of neuronal responses to drugs of abuse as well as drug related behaviors (Hitzemann *et al*. 2020; Ma *et al*. 2021; Ramos *et al*. 2021). Neuronal primary cilia offer new horizons for understanding how drugs of abuse affect the brain and how the neural basis of addiction develops, as well as unique potential therapeutic targets.

Primary cilia are specialized microtubule-based organelles formed during the mitotic process, resorbed, and then formed again following differentiation (Fuchs and Schwark 2004). In the adult brain, cilia remain relatively stable while exhibiting some dynamic structural responses to extracellular signals (Chakravarthy *et al*. 2012; Guadiana *et al*. 2016; Schou *et al*. 2015; Wheway *et al*. 2018). While much remains to be discovered about the capacities of primary cilia in the nervous system, they are implicated in metabolism, cognition/memory, and motivated behaviors (Kirschen and Xiong 2017; Ramos *et al*. 2021; Sherafat-Kazemzadeh *et al*. 2013; Vien *et al*. 2023). Disruption of ciliary function leads to phenotypes including obesity, cognitive impairments, disrupted food seeking, and impaired social motivation (Brinckman *et al*. 2013; Green and Mykytyn 2010; Poretti and Gerner 2016). Primary cilia are important for neuronal development, including proliferation, migration, differentiation and plasticity (Guemez-Gamboa *et al*. 2014; Higginbotham *et al*. 2013; Louvi and Grove 2011). Interference with ciliary signaling disrupts axon targeting, dendritic arborization, and the formation and maintenance of synapses (Guadiana *et al*. 2013; Guo *et al*. 2017; Guo *et al*. 2019; Lee and Gleeson 2010). Recent data also show that the PK2DL1 ion channel localizes to primary cilia and contributes to regulation of action potentials in interneurons, directly linking this organelle to neuronal excitability and function (Vien *et al*. 2023)

In order to carry out their cellular roles, primary cilia must be able to sense signals from other cells and physiological systems. Many of the neurobehavioral functions of primary cilia have been found to be mediated by ciliary reception of extracellular cues via a variety of GPCRs (Hilgendorf *et al*. 2016; Schou *et al*. 2015). Particularly relevant to the current study, multiple GPCRs known to be enriched in neuronal primary cilia also mediate drug responses and addiction-related behaviors. These include the serotonin 5HT-6 receptor (Brodsky *et al*. 2016; Eskenazi and Neumaier 2011; Ferguson *et al*. 2008; Frantz *et al*. 2002; Valentini *et al*. 2013; van Gaalen *et al*. 2010), the orphan GPCR, GPR88 (Ben Hamida *et al*. 2022; Ben Hamida *et al*. 2018; Jin *et al*. 2018; Laboute *et al*. 2020; Meirsman *et al*. 2019), and the melanin-concentrating hormone receptor MCHR1 (Berbari *et al*. 2008; Chee *et al*. 2019; Chung *et al*. 2009; Sun *et al*. 2013; Tyhon *et al*. 2006). In addition, the dopamine D1 receptor is capable of trafficking in and out of cilia, while activity at dopamine D2 receptor signaling can regulate cilia length (Domire *et al*. 2011; Miyoshi *et al*. 2014; Stubbs *et al*. 2022a; Stubbs *et al*. 2022b). It is still largely unknown what role the cilium itself plays in facilitating receptor-mediated drug responses, other than being a specialized region of the cell membrane where particular receptors are enriched.

We previously established a model for testing the neurobehavioral impact of targeting the intraflagellar-transport 88 (IFT88) gene for cilia removal on either GAD2-GABAergic or dopaminergic neurons (Ramos *et al*. 2021). Our findings indicated that neuronal cilia mediate locomotor responses to amphetamine in a cell-specific manner, which prompted further investigation of the role of neuronal cilia in behavioral responses to drugs of abuse. The current study expands upon our previous work by investigating whether cilia ablation produced similar effects on the response to cocaine. Cocaine and amphetamine are both considered psychostimulants but they are pharmacodynamically (Calipari *et al*. 2015; Kahlig and Galli 2003; Sulzer 2011) and pharmacokinetically distinct (Chiu and Schenk 2012; Mantle *et al*. 1976; Miller *et al*. 1980). Given the distinct pharmacological profiles of cocaine and amphetamine, as well as the diverse nature and range of the ciliary receptor signaling landscape, it is reasonable to expect that cilia loss may differentially regulate drug responses and warrants an investigation into each.

While locomotor responses have long been used as a proxy for measuring responses to psychostimulants and offer measures of drug induced neuroplasticity (Robinson and Berridge 2008; Segal and Mandell 1974; Vanderschuren and Kalivas 2000), the locomotor paradigm does not offer direct insight into the propensity of psychostimulants to produce brain reward. Conditioned place preference (CPP) is a commonly used paradigm within which the rewarding properties of drugs of all classes can be assessed (McKendrick and Graziane 2020; Schechter and Calcagnetti 1993; Tzschentke 2007). As multiple ciliary enriched receptors have been shown to modulate drug reward, a factor which contributes to abuse liability, we also increased the scope of our investigation by including a test of the effects of cell-specific cilia ablation on the rewarding properties of cocaine using a CPP paradigm.

## Materials and Methods

### Animals

Generation of neuron-specific cilia knockouts used for these experiments was previously detailed (Ramos *et al*. 2021). Briefly, for cilia loss on GABAergic neurons, Gad2IresCre (Jax stock 010802) mice were crossed with *Ift88* floxed mice, such that Gad2^+/c^:Ift88^+/F^ or Gad2^+/c^:Ift88^+/+^ mice are phenotypically wild-type (WT), and referred to as Gad2:Ift88^WT^, while Gad2^+/c^:Ift88^F/F^ mice are cilia knockouts and referred to as Gad2:Ift88^KO^ (Haycraft *et al*. 2007; Taniguchi *et al*. 2011). For cilia loss on dopaminergic neurons, DatIresCre mice (Jax stock, 006660, (Backman *et al*. 2006)) were used so that wildtype mice are Dat^+/c^:Ift88^+/F^ or Dat^+/c^:Ift88^+/+^ (Dat:Ift88^WT^) and knockout mice are DAT^+/c^:Ift88^F/F^ (Dat:Ift88^KO^). Breeding strategies and offspring genotypes are shown in Table 1. Mice were genotyped by extracting DNA from tail clippings with Extracta DNA Prep for PCR—Tissue (Quanta Biosciences) and specified products amplified using either GoTaq Green Mastermix (Promega) or 2x KAPA buffer. Mice of both sexes between 10 and 12 weeks of age were used for all experiments and were group housed 3–5 per cage from weaning until use. The estrous cycle of female mice was not monitored for these experiments (Meziane et al. 2007). Mice were maintained on a 12/12 light/dark cycle (lights on at 0700) with ad libitum access to food and water. Experiments were performed between 0800 and 1700. All procedures were approved by the University of Florida Institutional Animal Care and Use Committee and followed NIH guidelines.

### Locomotion

Locomotor activity was assessed in activity monitoring chambers (ENV-510) equipped with infrared beams to detect movement (MED-OFAS-MSU) (Med Associates, INC, Fairfax, VT). The chambers were housed within sound-attenuating cubicles, and locomotor activity was assessed in the dark (lights off in the cubicles). To assess the acute locomotor effects of cocaine, mice were first placed in the chambers and monitored for 1hr, followed by an i.p. injection of cocaine and monitoring for an additional hour. Activity in the chambers was determined using Med Associates software and calculated in 5 min bins. The two primary measures of interest were distance traveled (calculated from the Euclidean distance of all ambulatory episodes and expressed in centimeters) and repeated beam breaks (a measure of stereotypic behavior, calculated as repeated breaks of the same infrared beam). To assess locomotor sensitization, cocaine injections were given for an additional 4 days. On days 2-5, the pre-injection monitoring period was omitted and mice were injected and placed in the chambers for 1hr.

### Conditioned place preference

Conditioned place preference was assessed under a 21-day protocol, using a 3-chamber place preference apparatus equipped with infrared beams to monitor activity (ENV-3013, Med Associates, Inc). Day 1 consisted of a baseline test in which mice were placed in the central chamber for 5 minutes before being given access to the entire apparatus. Baseline preference for the two side chambers was assessed, and a biased subject assignment procedure was used such that cocaine was paired with the initially less-preferred chamber (Cunningham *et al*. 2006). On days 2-9 mice underwent place preference training wherein on alternating days they were injected and placed in either the saline-paired chamber or the cocaine-paired chamber. On day 10, for the initial CPP test, mice were given full access to the apparatus once again and activity was monitored to assess time spent in the saline-paired chamber vs. in the cocaine-paired chamber. On days 11-20 mice went through an extinction period in which they were given access to the full apparatus each day under the same conditions as the initial CPP test. Finally, on day 21, a reinstatement test was performed in which the mice were first administered cocaine at the same dose given during training before being given access to the full apparatus.

### Drug

Cocaine HCl (NIDA Drug Supply Program) was dissolved in sterile 0.9% saline vehicle, and administered via i.p. injection at a volume of 10.0 ml/kg prior to behavioral testing. The cocaine doses used (3.0, 10.0, 30.0 mg/kg) were chosen on the basis of published data showing that they bracket the range in which both acute locomotor stimulation and locomotor sensitization are observed (Collins et al. 2015)

### Data analyses

Statistical analyses were conducted in SPSS 29.0. The effects of acute cocaine were assessed separately in each strain of mice via multi-factor repeated measures ANOVA, with time bin as a within-subjects factor, and sex, dose, and genotype as between-subjects factors. Effects of sex and genotype on baseline locomotion (prior to cocaine injections) were assessed in a similar manner. The effects of repeated cocaine were also assessed separately in each strain using repeated measures ANOVA, with day as a within-subjects factor, and sex, dose, and genotype as between-subjects factors. Data from the CPP task were analyzed via two-(baseline preference, initial conditioned place preference, and reinstatement test) or three-(extinction) factor ANOVA. A full report of all statistical results for comparisons of locomotor activity and CPP performance is provided in **(Supplemental Table1)**. Greenhouse-Geisser corrections were used to correct for violations of sphericity in ANOVAs. For all analyses, p-values less than or equal to 0.05 were considered statistically significant.

## Results

### Effects of cilia ablation on activity in response to acute and repeated cocaine administration

To examine the effects of cilia ablation on activity in response to both acute and repeated cocaine, we used a 5-day administration regimen at doses shown previously to both acutely stimulate locomotion and induce sensitization to the locomotor stimulant effects of the drug (Collins et al. 2015). Day 1 of this regimen consisted of 1hr of monitoring of baseline activity, followed by injection of cocaine (or saline vehicle) and another hour of activity monitoring. On days 2-5, mice received injections immediately prior to 1hr of activity monitoring.

#### Effects of cilia loss genotypes on baseline activity

To evaluate the effects of cilia ablation on baseline activity, data from all mice prior to the first injections of cocaine (Day 1, collapsed across the 0, 3.0, 10.0, and 30.0 mg/kg doses) were compared between genotypes. In Dat:Ift88^KO^ mice, a three-factor ANOVA (genotype x time bin x sex) conducted on the distance traveled measure revealed a main effect of time bin, such that all mice decreased their distance traveled across bins (F(11,1375)=189.30, p<.001, Supplemental table 1), but no main effects or interactions involving genotype or sex (Fs<1.17, ps>.32) (Figure 1A, B). The same analysis conducted on the repeated beam break measure also revealed a reduction across time bins (F(11,1375)=150.14, p<.001; Figure 1C,D); in contrast to the distance traveled measure, however, there was a time bin x sex interaction, such that the reduction was initially more rapid in females than males (F(11,1375)=3.97, p=.003). Although repeated beam breaks were not significantly different between genotypes (main effect of genotype: F(1,125)=1.06, p=.30; genotype x time bin interaction: F(11,1375)=0.31, p=.88), there was a sex x genotype interaction, such that Dat:Ift88^KO^ mice showed more repeated beam breaks than WT in males but not in females (F(1,125)=4.07, p=.046; data not shown).

**Figure 1.**
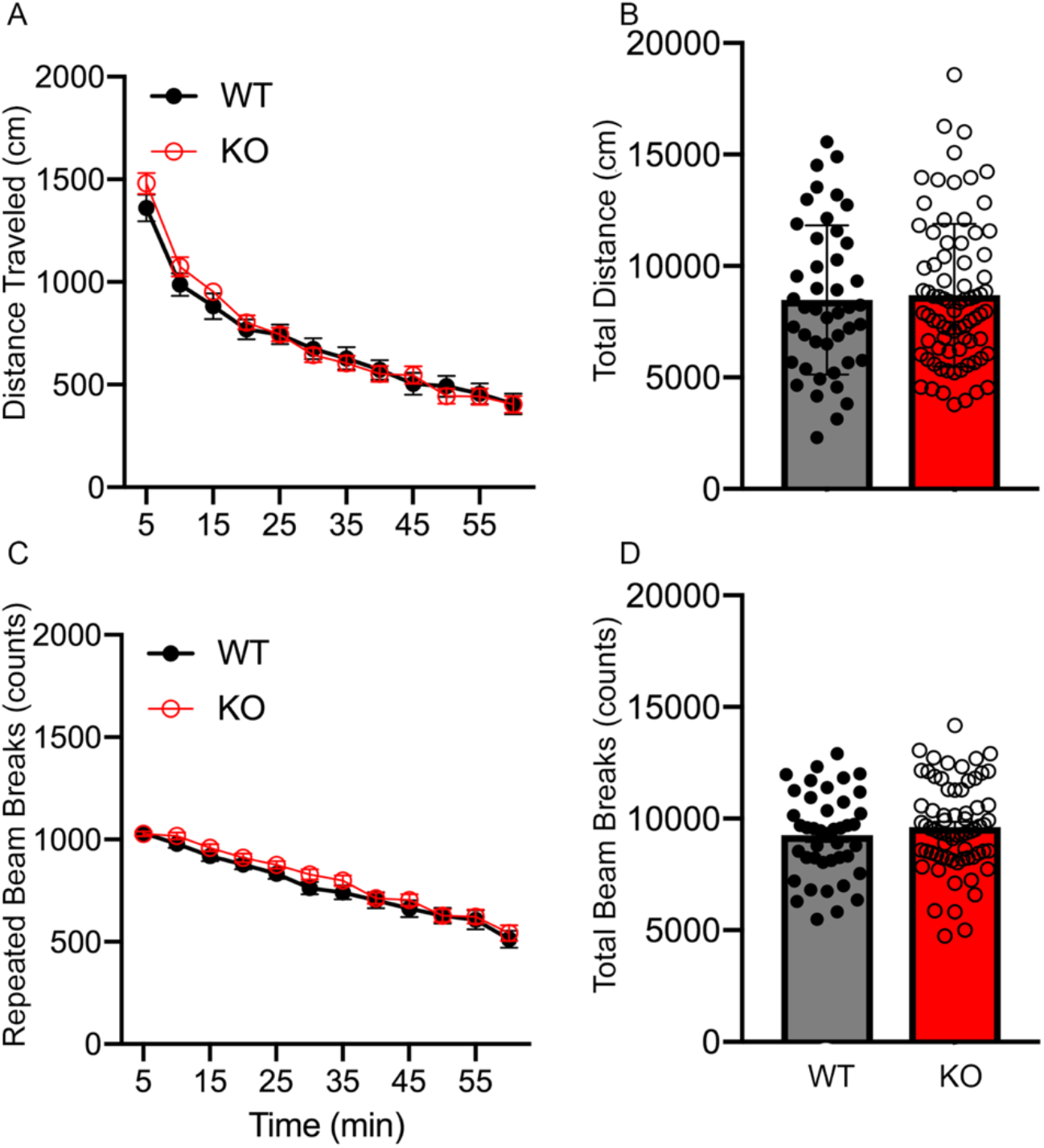
Dat:Ift88^KO^ mice do not show changes in basal activity. (A) Time course of horizontal distance traveled per 5 min bin of all WT & KO mice during an initial 1hr recording period. (B) Summed distance traveled in 1hr. (C) Time course of repeated beam breaks per 5 min bin of all WT & KO mice during an initial 1hr recording period. (D) Summed repeated beam breaks in 1hr.

In Gad2:Ift88^KO^ mice, analysis of distance traveled revealed a main effect of time bin, such that all mice decreased their distance traveled across bins (F(11,1067)=246.22, p<.001, Supplemental table 1), but no main effects or interactions involving sex or genotype (Fs<1.69, ps>.19, Figure 2A, B). The same analysis conducted on repeated beam breaks also revealed a decrease across bins (F(11,1067)=111.30, p<.001), but there were no main effects or interactions involving either sex or genotype (Fs<3.75, ps>.06, Figure 2C, D).

**Figure 2.**
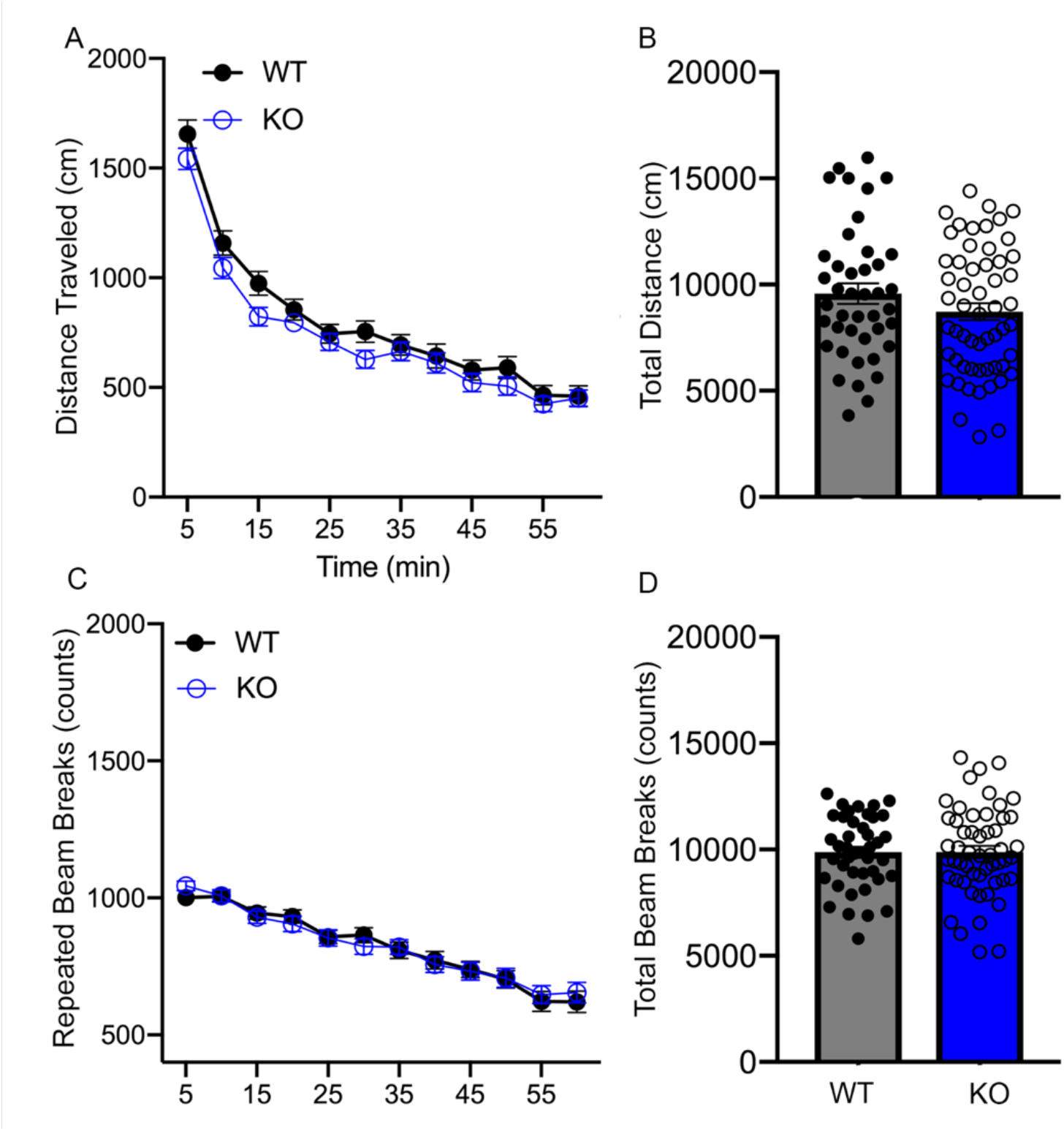
Gad2:Ift88^KO^ mice do not show changes in basal activity. (A) Time course of horizontal distance traveled per 5 min bin of all WT & KO mice during an initial 1hr recording period. (B) Summed distance traveled in 1hr. (C) Time course of repeated beam breaks per 5 min bin of all WT & KO mice during an initial 1hr recording period. (D) Summed repeated beam breaks in 1hr.

#### Effects of Dat:Ift88^KO^on acute cocaine-induced activity

To evaluate the effects of cocaine on activity in Dat:Ift88^KO^ mice, a 4-factor ANOVA (sex x dose X genotype X time bin) was conducted on data from the 60 minutes following injections (0, 3.0, 10.0, 30.0 mg/kg) (Figure 3A-D, Supplemental Table 2). On distance traveled, there was a main effect of dose (F(3,113)=180.72, p<.001) and time bin (F(11,1243)=60.84, p<.001) as well as a dose X time bin interaction (F(11,1243)=13.48, p<.001), such that distance was greater at higher doses, particularly in earlier time bins. There was also a main effect of sex (F(1,113)=5.57, p=.02), as well as a time bin x sex x dose (F(33,1243)=2.72, p=.002) interaction, such that females were more active than males, particularly in earlier time bins at the 30 mg/kg dose. There was also a genotype x sex x dose interaction (F(3,113)=3.62, p=.02), such that the effects of cocaine in KO and WT mice differed by sex (Supplemental Figure 1). To further explore the effects of genotype at each dose, follow-up ANOVAs (genotype x sex x time bin) were conducted on data from each dose condition. There were no main effects or interactions involving genotype following the 0, 3, or 30 mg/kg doses (Fs<1.61, ps>.14); at the 10 mg/kg dose, however there was a genotype x sex interaction (F(1,30)=7.24, p=.01), such that in females, distance traveled was less in KO compared to WT mice in females, whereas it was greater in KO compared to WT mice in males (Supplemental Figure 1A, C).

**Figure 3.**
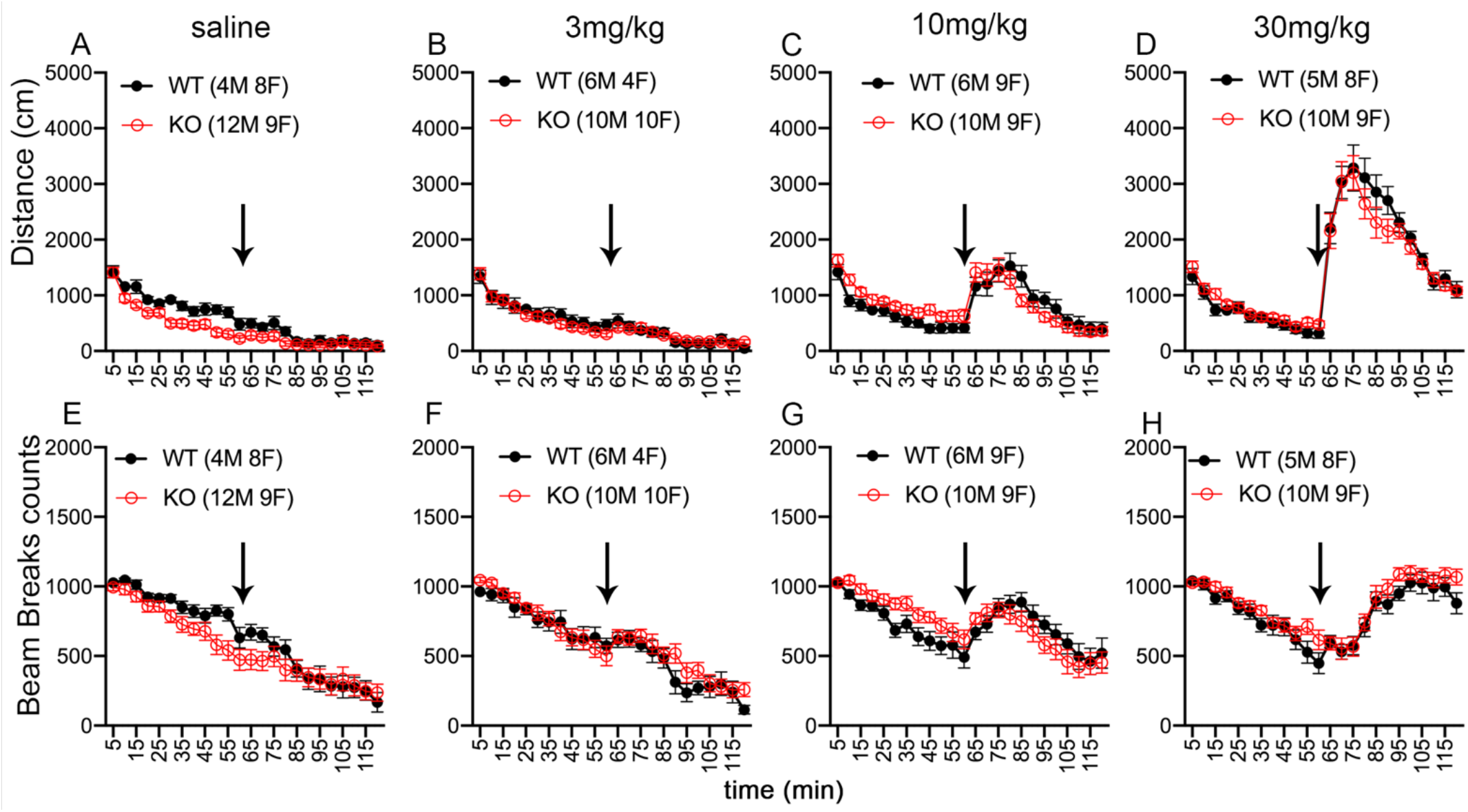
Dat:Ift88^KO^ effects on acute locomotor responses to cocaine. (A-D) Time course of horizontal distance traveled per 5 min bin in male & female WT & KO mice in response to A) Saline B) 3 mg/kg, C) 10 mg/kg, or D) 30 mg/kg. There were no main effects of genotype, but there was a genotype x sex x dose interaction (F(3,113)=3.62, p=.02), such that at the 10 mg/kg dose, there was a genotype x sex interaction (F(1,30)=7.24, p=.01). (E-H) Time course of repeated beam break per 5 min bin in male & female WT & KO mice in response to E) Saline F) 3 mg/kg, G) 10 mg/kg, or H) 30 mg/kg. No other significant differences related to genotype were observed. Arrows indicate time of IP administration of drug.

A comparable 4-factor ANOVA (sex x dose x genotype x time bin) was also conducted on the repeated beam break measure (Figure 3E-H, Supplemental Table 2). As with the distance traveled measure, this analysis revealed main effects of dose (F(3,113)=36.13, p<.001) and time bin (F(11,1243)=13.90, p<.001), as well as time bin x sex (F(11,1243)=2.31, p=.04) and time bin x dose (F(33,1243)=28.75, p<.001) interactions, but none of the main effects or interactions involving genotype reached statistical significance (Fs<3.00, ps>.05). In summary, there appeared to be limited effects of cilia loss on dopaminergic neurons on the acute locomotor response to cocaine, which were only evident on some measures at specific doses.

#### Effects of Gad2:Ift88^KO^on acute cocaine-induced activity

To evaluate the effects of cocaine on activity in Gad2:Ift88^KO^ mice, a 4-factor ANOVA (sex x dose x genotype x time bin) was conducted on data from the 60 minutes following injections (0, 3.0, 10.0, 30.0 mg/kg) (Figure 4A-D, Supplemental Table 2). On the distance traveled measure, there was a main effect of dose (F(3,85)=82.88, p<.001) and time bin (F(11,935)=56.81, p<.001) as well as a dose x time bin interaction (F(33,935)=12.46, p<.001), such that distance was greater at higher doses, particularly in earlier time bins. In contrast to the Dat:Ift88^KO^ mice, however, none of the other main effects or interactions on this measure reached statistical significance (Fs<3.13, ps>.08). A comparable 4-factor ANOVA (sex x dose x genotype x time bin) was also conducted on the repeated beam break measure (Figure 4E-H, Supplemental Table 2). As with the distance traveled measure in these mice, there was a main effect of dose (F(3,85)=34.33, p<.001) and time bin (F(11,935)=16.73, p<.001), as well as a time bin x dose interaction (F(33,935)=10.93, p<.001), such that differences in stereotypy between doses were more pronounced at later time bins. There was also a time bin x sex x dose interaction (F(33,935)=1.64, p=.048), but none of the main effects or interactions involving genotype reached statistical significance (Fs<1.28, ps>.27). In summary, there appeared to be no obvious effects of cilia loss on GABAergic neurons on the acute locomotor response to cocaine.

**Figure 4.**
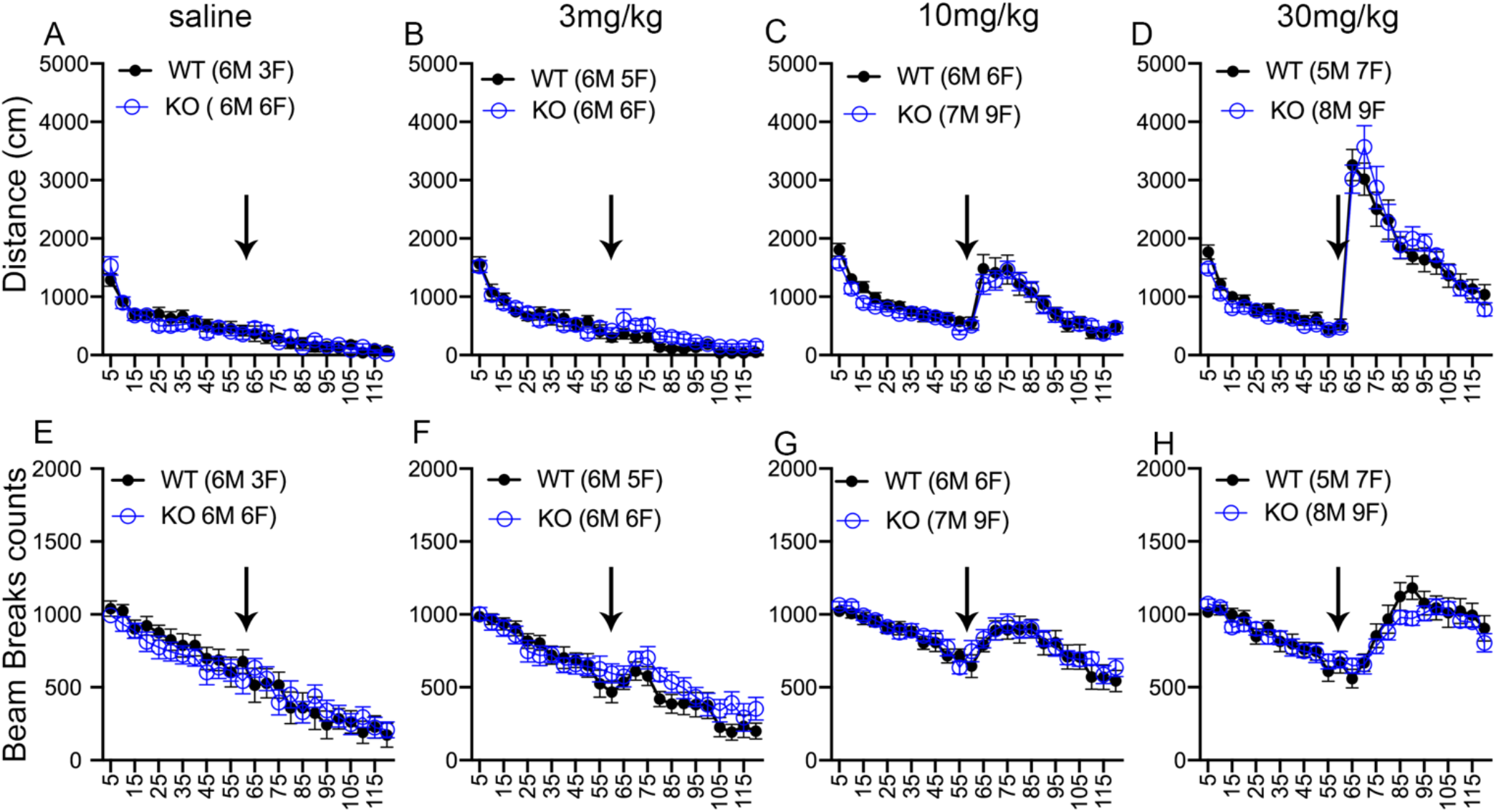
Gad2:Ift88^KO^ effects on acute locomotor responses to cocaine. (A-D) Time course of horizontal distance traveled per 5 min bin in male & female WT & KO mice in response to A) Saline B) 3 mg/kg, C) 10 mg/kg, or D) 30 mg/kg. (E-H) Time course of repeated beam break per 5 min bin in male & female WT & KO mice in response to E) Saline F) 3 mg/kg, G) 10 mg/kg, or H) 30 mg/kg. No significant differences related to genotype were observed. Arrows indicate time of IP administration of drug.

#### Effects of Dat:Ift88^KO^on plasticity of cocaine-induced activity

To determine whether cilia ablation on DAT expressing neurons affects the plasticity of the locomotor response to repeated cocaine administration (i.e., locomotor sensitization), the distance traveled and repeated beam break measures were evaluated across 5 days of cocaine or vehicle administration. For these analyses, activity on each day was collapsed across the 12 time bins and analyzed as total distance/beam breaks in 1hr. On the distance traveled measure, a 4-factor ANOVA (day x genotype x sex x dose) revealed main effects of day (F(4,452)=72.91, p<.001) and dose (F(3,113)=186.10, p<.001), as well as a day x dose interaction (F(12,452)=8.34, p<.001), such that the increase in distance across days was greater at higher doses (Figure 5A-D, Supplemental Table 3). There was also a main effect of genotype (F(1,113)=6.61, p=.01) as well as dose x genotype (F(3,113)=4.96, p=.003) and dose x day x genotype (F(12,452)=2.27, p=.02) interactions, such that the cocaine-induced increase in distance traveled across days differed between WT and KO mice. To further explore the effects of genotype at each dose, follow-up ANOVAs (genotype x sex x day) were conducted on data from each dose. These analyses revealed no main effects or interactions involving genotype in the 0 mg/kg group, but significant effects of genotype in the 3.0, 10.0, and 30.0 mg/kg groups. In the 3 mg/kg group, there was a main effect of genotype (F(1,26)=6.79, p=.02), such that KO mice traveled a greater distance than WT mice (Figure 5B). Surprisingly, this effect was reversed at the 10 mg/kg dose, at which there was a sex x genotype interaction (F(1,30)=4.27, p=.048), such that KO mice traveled a shorter distance than WT, but only in females (Figure 5C, Supplemental Figure 2B, E). KO mice also traveled a shorter distance than WT at the 30 mg/kg dose (main effect of genotype, F(1,28)=9.25, p=.005; day x genotype interaction, F(4,112)=2.76, p=.04), but this effect was independent of sex (Figure 5D, Supplemental Figure 3C, F)

**Figure 5.**
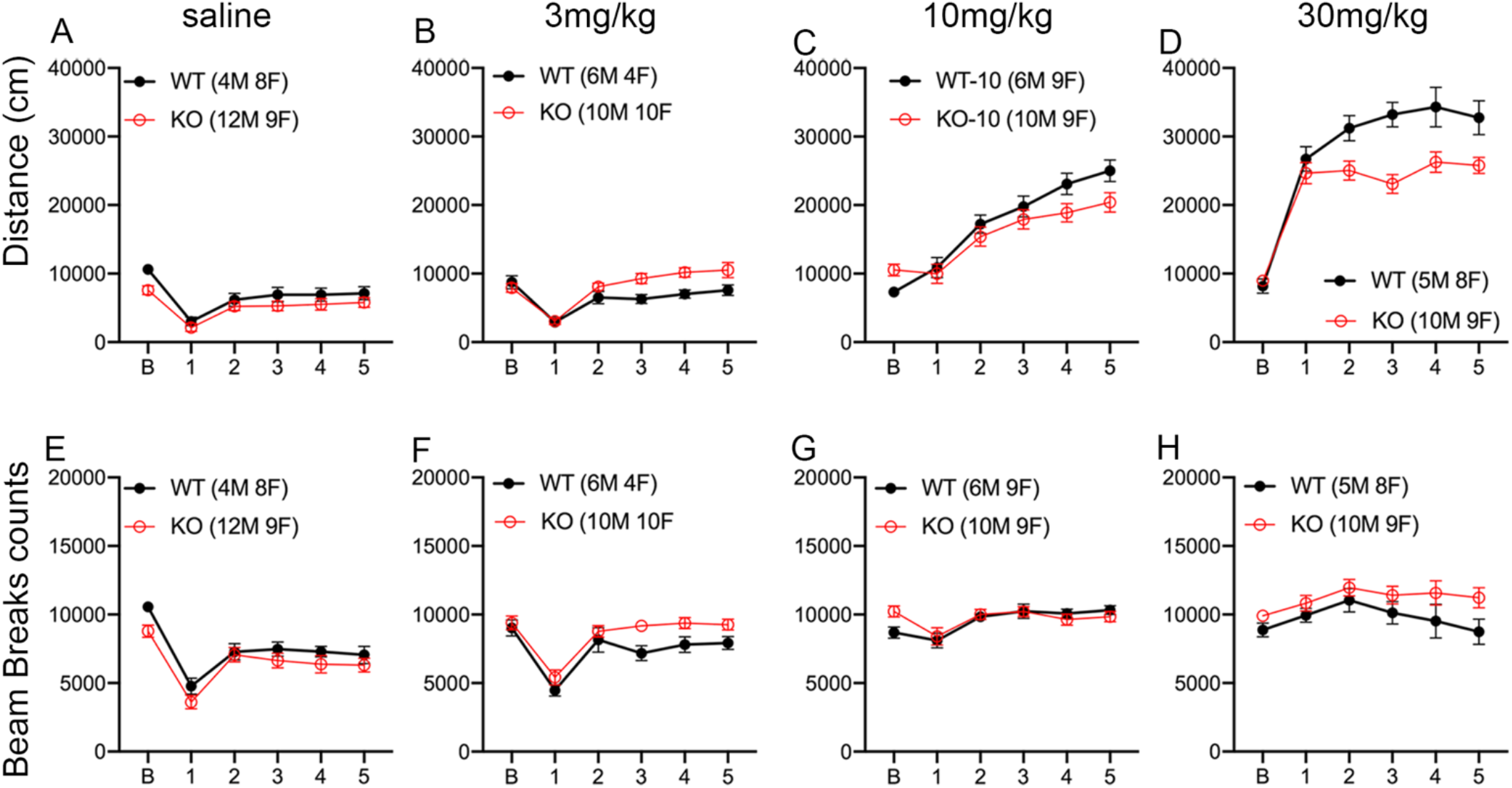
Dat:Ift88^KO^ alters cocaine-induced locomotor plasticity. (A-D) Total distance traveled in 1hr following 5 daily injections in male & female WT & KO mice in response to A) Saline B) 3 mg/kg, C) 10 mg/kg, or D) 30 mg/kg. Dat:Ift88^KO^ showed significant differences in locomotor sensitization (4-way ANOVA, main effect of genotype, F_(1, 113)_ =6.61, p=.01). KO mice displayed enhanced cocaine-induced locomotion across the 5 days at 3mg/kg IP (F(1,26)=6.79, p=.02) but reduced sensitization to the locomotor stimulant effects at 10/mg/kg (sex x genotype interaction, F(1,30)=4.27, p=.048) and 30mg/kg (main effect of genotype, F(1,28)=9.25, p=.005 (E-H) Total repeated beam breaks in 1hr following 5 daily injections in in male & female WT & KO mice in response to E) Saline F) 3 mg/kg, G) 10 mg/kg, or H) 30 mg/kg. No significant differences related to a main effect of genotype were observed, although there was a sex x genotype interaction (F(1,113)=4.28, p=.04).

A comparable 4-factor ANOVA (sex x dose x genotype x day) was also conducted on the repeated beam break measure (Figure 5E-H, Supplemental Table 3). As with the distance traveled measure, this analysis revealed main effects of dose (F(3,113)=26.21, p<.001) and day (F(4,452)=51.57, p<.001), as well as a dose x day interaction (F(12,452)=6.01, p<.001), such that the change in stereotypy across days was greater at some doses than others (Figure 5E-H). In addition, there was a sex x genotype interaction (F(1,113)=4.28, p=.04), such that in males, KO mice showed greater stereotypy than WT, whereas there was no genotype difference in females (Supplemental Figure 3). Considering all of these data together, it appears that cilia ablation on DAT expressing neurons results in dose-dependent alterations in sensitization to the locomotor stimulant effects of cocaine.

#### Effects of Gad2:Ift88^KO^ on plasticity of cocaine-induced activity

To determine whether cilia ablation on GAD2 expressing neurons affects the plasticity of the locomotor response to repeated cocaine administration, an analysis approach comparable to that performed on data from Dat:Ift88^KO^ mice was conducted. On the distance traveled measure, a 4-factor ANOVA (day x genotype x sex x dose) revealed main effects of day (F(4,340)=40.27, p<.001) and dose (F(3,85)=70.41, p<.001), as well as a day x dose interaction (F(12,340)=4.87, p<.001), such that the increase in distance across days was greater at higher doses (Figure 6A-C, Supplemental Table 3). In contrast to the Dat:Ift88^KO^ mice, however, there were no main effects or interactions involving genotype (Fs<1.73, ps>.16), although there was a main effect of sex, such that males traveled a greater distance than females. Analyses of the repeated beam break measure revealed results similar to those with the distance traveled measure, with a main effect of day (F(4,340)=31.27, p<.001) and dose (F(3,85)=20.68, p<.001), as well as a day x dose interaction (F(12,340)=6.00, p<.001)(Figure 6D-F, Supplemental Table 3). As with distance traveled, there was also a main effect of sex (F(1,85)=5.11, p=.03), with males showing greater stereotypy than females. In summary, there appeared to be no effects of cilia ablation from GAD2 expressing neurons on sensitization to the locomotor stimulant effects of cocaine.

**Figure 6.**
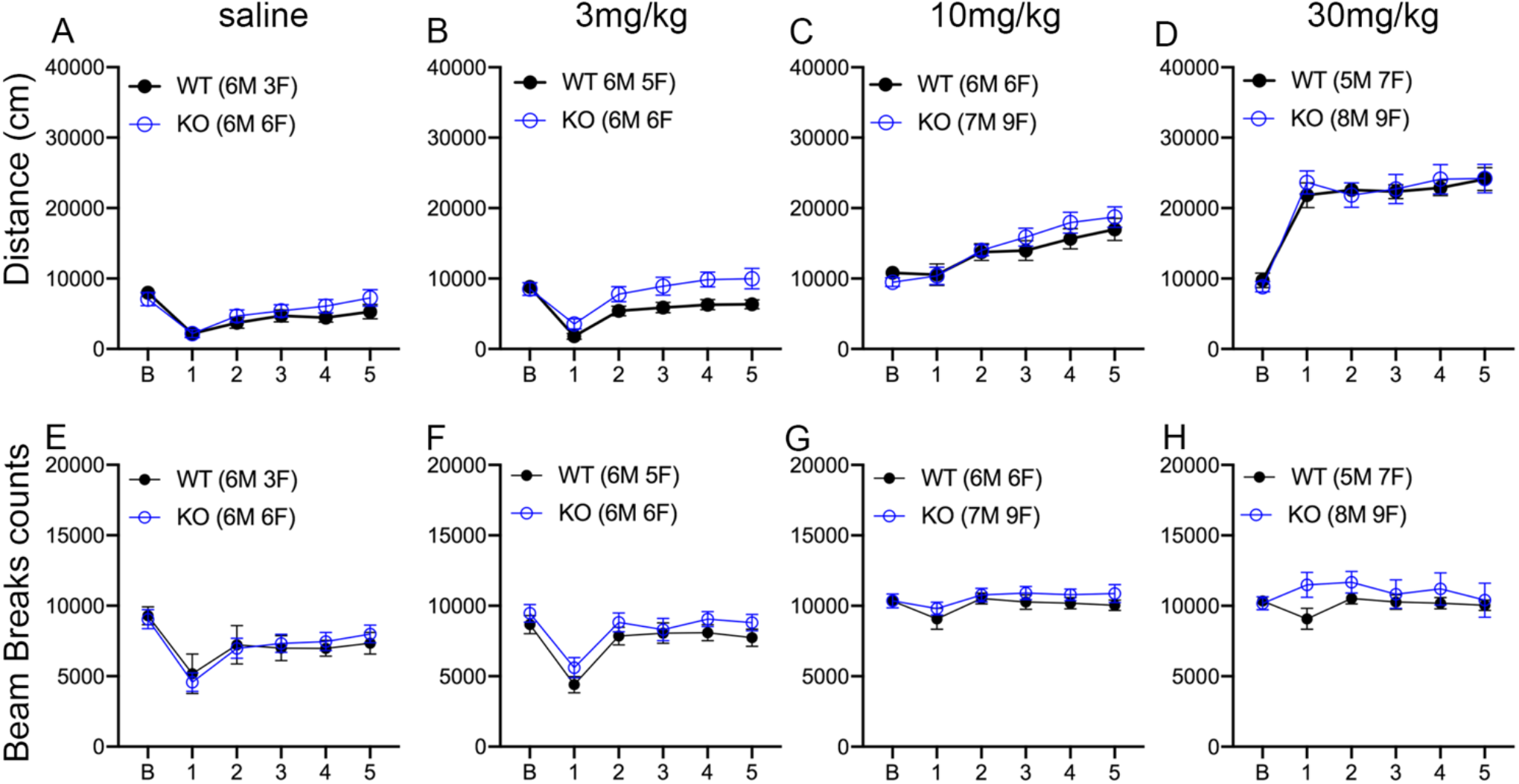
Gad2:Ift88^KO^ does not significantly alter cocaine-induced locomotor plasticity. (A-D) Total distance traveled in 1hr following 5 daily injections in male & female WT & KO mice in response to A) Saline B) 3 mg/kg, C) 10 mg/kg, or D) 30 mg/kg. No other significant differences due to genotype were observed. (E-H) Total repeated beam breaks in 1hr following 5 daily injections in in male & female WT & KO mice in response to E) Saline F) 3 mg/kg, G) 10 mg/kg, or H) 30 mg/kg. As with distance traveled significant differences due to genotype were not detected.

### Effects of cilia ablation on cocaine conditioned place preference

To evaluate the effects of cilia ablation on DAT- and GAD2-expressing neurons on the rewarding effects of cocaine, Dat:Ift88^KO^ and Gad2:Ift88^KO^ mice were trained and tested for cocaine CPP. A dose of 10.0 mg/kg cocaine was initially used for both mouse lines, but preliminary experiments showed that both KO and WT mice of the DAT:Ift88 line failed to show robust CPP, and thus a 30 mg/kg dose was used for this line.

#### Effects of Dat:Ift88KO on cocaine conditioned place preference

There were no differences between groups during initial baseline preference testing (two-factor ANOVA, genotype x sex, Fs<0.84, ps>.38). Following 8 days of pairing (alternating coke and saline injections), mice underwent a 15-minute place preference test, in which the time spent on the cocaine-paired vs. the saline-paired side of the apparatus was measured, and a ratio between the times on each side was calculated (paired/unpaired), with values greater than 1 representing preference for the paired side. A two-factor repeated measures ANOVA (genotype x sex) revealed a main effect of genotype (F(1,11)=8.39, p=.02), such that preference for the paired side was attenuated in KO relative to WT mice (Figure 7A), but no main effect of sex or sex x genotype interaction (Supplemental table 4). Following this initial test, mice underwent 10 days of extinction sessions, each of which was identical to the initial place preference test. During these sessions, a three-factor repeated measures ANOVA (extinction day x genotype x sex) revealed no main effects or interactions involving genotype (Figure 7B, Supplemental Table 4). In contrast, during the reinstatement test (a 15 min session following injection of 30 mg/kg cocaine) conducted on the day following the last extinction session, there was a main effect of genotype (F(1,11)=6.00, p=.03), such that, as in the initial test, preference for the cocaine-paired side was attenuated in KO relative to WT mice (Figure 7A). Considered together, these data show that ablation of cilia on dopaminergic neurons attenuates learning about/expression of the rewarding/reinforcing effects of cocaine, but does not affect extinction of such behavior.

**Figure 7.**
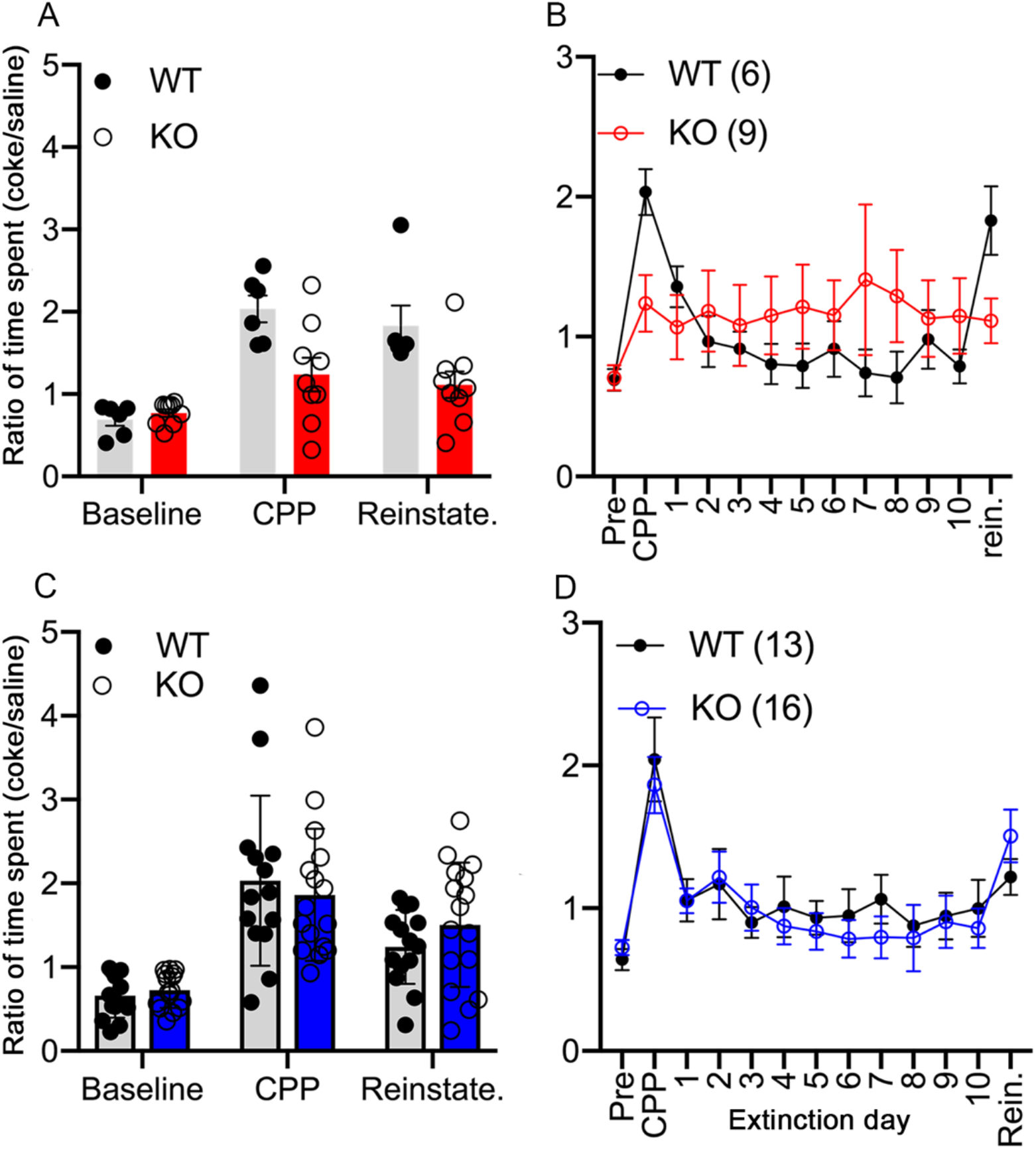
Effects of cell-specific cilia ablation on the rewarding properties of cocaine. (A) Dat:Ift88^ko^ mice show a reduction in time spent in the cocaine paired chamber in both the CPP and reinstatement tests (two-factor repeated measures ANOVA, CPP main effect of genotype (F(1,11)=8.39, p=.02), Reinstate. main effect of genotype (F(1,11)=6.00, p=.03)). (B) Ratio of time spent in cocaine paired side for each day including extinction days, a three-factor repeated measures ANOVA (extinction day x genotype x sex) revealed no main effects or interactions involving genotype. (C) Gad2:Ift88^ko^ mice showed no changes in time spent in the cocaine paired chamber in both the CPP and reinstatement tests (two-factor repeated measures ANOVA, main effect of genotype (F(1,25)=2.41, p=.13), Reinstate. main effect of genotype (F(1,11)=6.00, p=.03)). (D) A main effect of genotype emerged during the 10 extinction sessions (F(1,25)=4.22, p=.05), with KO mice showing attenuated preference for the cocaine-paired side across all sessions.

#### Effects of Gad2:Ift88^KO^on cocaine conditioned place preference

During initial baseline preference testing, there was a significant sex difference, such that, across genotypes, males showed greater preference for the side to-be-paired with cocaine compared to females (F(1,11)=5.30, p=.03), but no effects of genotype (Figure 7C, supplemental table 4). This sex difference persisted during the initial place preference test (F(1,25)=9.99, p<.001), although there were still no effects of genotype (Supplemental Table 4). In contrast, a main effect of genotype emerged during the 10 extinction sessions (F(1,25)=4.22, p=.05), with KO mice showing attenuated preference for the cocaine-paired side across all sessions, though there were no main effects or interactions involving sex. This difference was absent during the reinstatement session, in which there were no effects of either genotype or sex (Fs<1.02, ps>.32). Considered together, these results suggest that although cilia ablation on GAD2 neurons alters the course of extinction, it does not appear to modulate cocaine reward/reinforcement.

## Discussion

Primary cilia remain an underexplored landscape in which to study drug responses and behaviors relevant to substance use. Several studies have now explored the role of neuronal ciliary-enriched GPCRs in modulating drug responses (Laboute *et al*. 2020; Ramos *et al*. 2021). In addition to receptor-based signaling, however, the ciliary structure itself may be an important feature. Therefore, our objective was to explore how the loss of the ciliary structure impacts drug responses using cell-specific knockout of the IFT88 gene. This current work expands on how cilia loss from specific populations of neurons impacts a variety of neurobehavioral phenotypes (Alhassen *et al*. 2023; Mustafa *et al*. 2019; Ramos *et al*. 2021; Stubbs *et al*. 2022b). Ultimately our findings support that the effects of cilia loss on drug responses are cell type- and drug-specific, as well as dose-dependent, and may differ between sexes.

Cilia are critical for development, and disruption of ciliary IFT genes during development, including IFT88, can produce severe developmental defects and limit survival (McIntyre *et al*. 2012; Rosenbaum and Witman 2002; Tian *et al*. 2017). We therefore used Cre/LoxP-mediated IFT88 removal to ablate cilia on specific neuron populations post-differentiation, and we previously reported that this leads to successful loss of cilia without producing noticeable changes in neuron survival at the ages used in this study (Ramos *et al*. 2021). Confirming cilia loss can be difficult in neural tissue; however, several studies have used the structural proteins acetylated α-tubulin (Coveney *et al*. 2022a; Coveney *et al*. 2022b; Green *et al*. 2018; Joiner *et al*. 2015; Lee *et al*. 2021; McDermott *et al*. 2010; Toomer *et al*. 2017), polyglutamylated α-tubulin (McIntyre *et al*. 2012), and de-tyrosinated tubulin (Palla *et al*. 2022) to confirm that IFT88 mutations lead to cilia loss on cell types throughout the body. Other studies have used ciliary-enriched transmembrane and membrane-associated proteins to verify cilia loss on neurons (Berbari *et al*. 2014; Bowie and Goetz 2020; Domire and Mykytyn 2009; Koemeter-Cox *et al*. 2014; Mustafa *et al*. 2019; Ramos *et al*. 2021; Sipos *et al*. 2018; Sterpka and Chen 2018; Stubbs *et al*. 2022b; Yang *et al*. 2022). Multiple lines of evidence support the conclusion that IFT88 disruption leads to cilia loss, and an increasing number of studies are using IFT-mediated cilia ablation to study the impacts of cilia loss on neurobehavioral outcomes (Alhassen *et al*. 2023; Amador-Arjona *et al*. 2011; Bowie and Goetz 2020; Chizhikov *et al*. 2007; Lee *et al*. 2022; Ramos *et al*. 2021; Rhee *et al*. 2016; Stubbs *et al*. 2022b; Tu *et al*. 2023; Yang *et al*. 2021).

Given the broad range of the ciliary signaling landscape, cilia ablation may alter physiological processes that alter drug responses outside of the loss of ciliary signaling pathways. In particular, disruption of ciliary function is associated with obesity and changes in metabolism (DeMars *et al*. 2023; Lee *et al*. 2022; Siljee *et al*. 2018; Stubbs *et al*. 2022b). We previously reported that GAD2-GABAergic neuron cilia ablation led to a significant decrease in body weight of about 15%, while dopaminergic neuron cilia ablation did not lead to differences in body weight (Ramos *et al*. 2021). Since cilia loss on either cell type did not alter baseline locomotor activity or responses to saline, it remains possible that changes in food consumption or increased metabolism may contribute to the effects of GABAergic cilia loss on locomotor responses to psychostimulants; however, increased metabolism typically leads to a need for higher doses of a drug to achieve an effect (Sharma and McNeill 2009; Susa *et al*. 2023). As Gad2:Ift88^KO^ mice displayed an increased locomotor response at the lowest cocaine dose, along with the lack of significant effects on locomotion at higher cocaine doses or effects on cocaine reward, this may suggest that metabolic changes are not responsible for the effects of GABAergic cilia loss on the response to psychostimulants.

We observed that primary cilia loss from GABAergic or dopaminergic neurons did not impact locomotor responses to acute administration of cocaine at a range of doses (3, 10, & 30mg/kg). This contrasts with our previous study in which cilia loss on either GAD2-gabaergic or dopaminergic neurons reduced acute locomotor responses to 3mg/kg amphetamine, a dose that produced levels of activity comparable to 30mg/kg cocaine (Ramos *et al*. 2021). Cocaine and amphetamine are pharmacologically distinct in multiple ways, which may contribute to variability in the effects of cilia loss on acute drug responses. For example, the half-life of amphetamine following i.p. administration is up to 4-fold longer than that of cocaine, and the two drugs have different affinities for monoamine transporters (Benuck *et al*. 1987; Fuller *et al*. 1972; Hicks *et al*. 1980; Howell and Kimmel 2008).

In contrast to acute responses, locomotor sensitization in Gad2:IFT88^KO^ mice was altered, though only at the lowest dose. GAD2 is expressed in both Drd1+ and Drd2+ medium spiny neurons in the rodent striatum (Ferhat *et al*. 2023). Activation of Drd1+ neurons is known to promote locomotion (Freeze *et al*. 2013; Kravitz *et al*. 2010), while Drd2+ neuron activation and DRD2 agonism can inhibit locomotion (Barrozo *et al*. 2015; Durieux *et al*. 2009; Ford 2014). Disruption of either cilia (through Ift88 knockout), or ciliary trafficking of GPCRs (via Bbs1 knockout), specifically on Drd1+ neurons leads to reductions in general locomotion (Stubbs *et al*. 2022b). Targeting of ciliary signaling on Drd1+ neurons through either approach also lead to increased bodyweight (Stubbs *et al*. 2022b). In our GAD2 cilia KO mice, general locomotion is not altered, and bodyweight is decreased. Since Gad2:Ift88^KO^ would lead to cilia loss on both Drd1+ and Drd2+ striatal neurons, disruption of ciliary dopamine signaling in both populations may have opposing effects, obscuring overall impacts on locomotion. Drd2 receptors have significantly higher affinity for dopamine than Drd1 receptors (Keeler *et al*. 2014; Martel and Gatti McArthur 2020). Disruption of Drdr2 signaling in mice has been shown to have dose-dependent effects such that Drd2 KO reduces the rewarding properties of cocaine, but only at a low dose of 2.5mg/kg (Welter *et al*. 2007). Thus, the impact of eliminating ciliary signaling from both Drd1+ and Drd2+ neurons on cocaine-induced locomotion may only be observable at low doses at which D2 receptors are primarily occupied, and the disruption of Drd2+ neuron function due to cilia loss may remove inhibition of locomotion, leading to the observed increase in cocaine response.

In this study we extend an analysis of cilia loss from locomotor behaviors to the rewarding effects of stimulants through the use of the CPP assay. We find that loss of cilia from GAD2 neurons has no effect on development of cocaine CPP or its reinstatement, although it did appear to enhance extinction of the CPP. In contrast, cilia loss from DAT neurons caused a significant reduction in both the development/expression and reinstatement of cocaine CPP. This effect is likely not due to differential sensitivity of these animals to the cocaine dose, as acute locomotor responses to the same dose were not different. Notably, the latter results are consistent with the reduced locomotor sensitization in Dat:Ift88^KO^ mice, and in combination, they suggest that loss of cilia on DAT neurons changes plasticity and impairs cocaine-induced CPP learning and/or expression.

Several ciliary-enriched receptors are known to modulate effects of psychostimulants. The melanin-concentrating hormone receptor (MCHR1) is enriched in primary cilia in multiple brain regions, including the striatum (Diniz *et al*. 2020; Jasso *et al*. 2021), and is known to modulate dopamine release and behavioral responses to cocaine and amphetamine (Chung *et al*. 2009; Geuzaine *et al*. 2014; Morganstern *et al*. 2020; Pissios *et al*. 2008; Tyhon *et al*. 2006). MCH is produced primarily in neurons in the lateral hypothalamus (LH), which project to regions throughout the brain including the striatum, nucleus accumbens, and cortex, where MCHR1 is expressed on GABAergic interneurons (Chee *et al*. 2013; Diniz and Bittencourt 2017). The evidence for how MCH signaling may alter neurobehavioral responses to psychostimulants is somewhat mixed. For example, MCH application has been shown to potentiate dopamine-induced neuronal firing in the striatum and increase cocaine-induced locomotion, suggesting that it may augment psychostimulant effects (Chung *et al*. 2009). In contrast, knockout of MCHR1 also increases dopamine release in response to the selective dopamine reuptake inhibitor GBR12909, and potentiates GBR- and amphetamine-induced locomotion, suggesting it may play an inhibitory role in this context (Chee *et al*. 2019). While it is unclear whether signaling pathways of ciliary-enriched receptors remain intact after cilia ablation, cilia loss may lead to elimination of ciliary receptor signaling pathways, including MCHR1. Yet, cilia loss from Gad2 neurons inhibited acute locomotor responses to amphetamine (largely in male mice) and enhanced some aspects of sensitization (Ramos *et al*. 2021), while only increasing cocaine’s effect on sensitization at a low dose (this study). Thus, it appears that other ciliary signaling pathways beyond MCHR1 may be disrupted by cilia removal.

The serotonin receptor HTR6 is enriched in primary cilia of medium spiny neurons in the striatum and differentially modulates responses to psychostimulants (Brodsky *et al*. 2017; Dupuy *et al*. 2023; Frantz *et al*. 2002; Sheu *et al*. 2022). High doses of amphetamine increase serotonin levels (Kuczenski and Segal 1989), whereas most doses of cocaine will alter serotonin signaling (Wei *et al*. 2018). HTR6 antagonism potentiates the dopamine releasing and locomotor effects of amphetamine, but has no significant effect on the same measures of cocaine responses (Frantz *et al*. 2002). While overexpression of striatal HTR6 increases operant responding for cocaine, it does not impact locomotor responses to the drug (Brodsky *et al*. 2016; Ferguson *et al*. 2008). These findings suggest that deficits in HTR6 signaling due to cilia loss could contribute to the reduction in amphetamine-induced locomotion caused by cilia ablation, while leaving acute cocaine responses largely unaffected.

Whereas MCHR1 and HTR6 show ciliary enrichment in the striatum, neither appear to be significantly enriched in cilia of midbrain dopaminergic neurons (Brailov *et al*. 2000; Diniz *et al*. 2020; Dupuy *et al*. 2023; Jasso *et al*. 2021). The impacts of dopaminergic cilia ablation on psychostimulant responses are potentially regulated by different mechanisms (Jasso *et al*. 2021; Roberts *et al*. 2002). It has previously been shown that cilia loss from DAT-positive neurons leads to a reduction in substantia nigra to striatum projections and a reduction in striatal dopamine (Mustafa *et al*. 2021). One explanation for reduced behavioral responses in both the current and previous study could be that cilia loss leads to reduced dopamine release upon stimulation with either amphetamine or cocaine. This therefore might suggest not a cilia-specific signaling defect *per se*, but a role for primary cilia in the complete maturation and maintenance of dopaminergic neurons.

In summary, we show that cilia loss on different populations of neurons has distinct effects on both cocaine-induced locomotor plasticity and development/expression of cocaine CPP in mice. We further show that cocaine-induced locomotor activity can differ between sexes following the same ciliary manipulation. Finally, in combination with our previous work, we show that the effects of stimulant type on locomotor activity of mice with cilia loss from the same population of neurons differs. This opens the possibility that ciliary signaling differently modulates responses to different psychostimulant drugs of abuse. Most broadly, these results provide further evidence that neuronal cilia play important, though underexplored, roles in reward signaling in the brain.

## Funding/Acknowledgements

This work was supported by NIH R21DA047623 to JCM and BS, and NIH R01DC019379 to JCM. We thank the Drug Supply Program at the National Institute on Drug Abuse for kindly providing cocaine HCl.

