## Supplemental Tables for "Cilia loss on distinct neuron populations differentially alters cocaine-induced locomotion and reward"

| **Baseline activity** |  |
| --- | --- |
| *Dat:Ift88 KO* |  |
| Locomotor activity |  |
| Genotype | F(1,125)=0.00, p=.99 |
| Sex | F(1,125)=0.01, p=.98 |
| Time Bin | F(11,1375)=189.30, p<.001 |
| Genotype x Sex | F(1,125)=0.85, p=.36 |
| Genotype x Time Bin | F(11,1375)=1.17, p=.32 |
| Sex x Time Bin | F(11,1375)=0.86 p=.52 |
| Genotype x Sex x Time Bin | F(11,1375)=0.50, p=.79 |
| Repeated Beam Breaks |  |
| Genotype | F(1,125)=1.06, p=.30 |
| Sex | F(1,125)=1.21, p=.27 |
| Time Bin | F(11,1375)=150.14, p<.001 |
| Genotype x Sex | F(1,125)=4.07, p=.046 |
| Genotype x Time Bin | F(11,1375)=0.31, p=.88 |
| Sex x Time Bin | F(11,1375)=3.97, p=.003 |
| Genotype x Sex x Time Bin | F(11,1375)=1.09, p=.36 |
| *Gad2:Ift88 KO* |  |
| Locomotor activity |  |
| Genotype | F(1,97)=0.91, p=.34 |
| Sex | F(1,97)=1.35, p=.25 |
| Time Bin | F(11,1067)=246.22, p<.001 |
| Genotype x Sex | F(1,97)=1.68, p=.20 |
| Genotype x Time Bin | F(11,1067)=1.39, p=.20 |
| Sex x Time Bin | F(11,1067)=1.13, p=.34 |
| Genotype x Sex x Time Bin | F(11,1067)=1.39, p=.20 |
| Repeated Beam Break |  |
| Genotype | F(1,97)=0.06, p=.80 |
| Sex | F(1,97)=3.74, p=.06 |
| Time Bin | F(11,1067)=111.30, p<.001 |
| Genotype x Sex | F(1,97)=1.17, p=.28 |
| Genotype x Time Bin | F(11,1067)=0.89, p=.51 |
| Sex x Time Bin | F(11,1067)=1.83, p=.08 |
| Genotype x Sex x Time Bin | F(11,1067)=1.03, p=.41 |

**Supplemental Table 1:** Statistical analysis of 1hr baseline locomotor activity and repeated beam breaks of all Dat:Ift88 and Gad2:Ift88 mice.

| **Acute cocaine** |  |
| --- | --- |
| *Dat:Ift88 KO* |  |
| Locomotor activity |  |
| Genotype | F(1,113)=0.84, p=.36 |
| Sex | F(1,113)=5.57, p=.02 |
| Dose | F(3,113)=180.72, p<.001 |
| Time Bin | F(11,1243)=60.84, p<.001 |
| Genotype x Sex | F(1,113)=1.04, p=.31 |
| Genotype x Dose | F(3,113)=0.21, p=.89 |
| Genotype x Time Bin | F(11,1243)=1.05, p=.38 |
| Sex x Dose | F(3,113)=1.50, p=.22 |
| Sex x Time Bin | F(11,1243)=2.29, p=.07 |
| Dose x Time Bin | F(33,1243)=13.48, p<.001 |
| Genotype x Sex x Dose | F(3,113)=3.62, p=.02 |
| Genotype x Sex x Time Bin | F(11,1243)=1.24, p=.29 |
| Genotype x Dose x Time Bin | F(33,1243)=0.79, p=.65 |
| Sex x Dose x Time Bin | F(33,1243)=2.72, p=.002 |
| Genotype x Sex x Dose x Time Bin | F(33,1243)=0.87, p=.57 |
| Repeated Beam Breaks |  |
| Genotype | F(1,113)=0.53, p=.82 |
| Sex | F(1,113)=0.24, p=.63 |
| Dose | F(3,113)=36.13, p<.001 |
| Time Bin | F(11,1243)=13.90, p<.001 |
| Genotype x Sex | F(1,113)=3.00, p=.09 |
| Genotype x Dose | F(3,113)=0.53, p=.67 |
| Genotype x Time Bin | F(11,1243)=1.26, p=.28 |
| Sex x Dose | F(3,113)=2.38, p=.07 |
| Sex x Time Bin | F(11,1243)=2.31, p=.04 |
| Dose x Time Bin | F(33,1243)=28.75, p<.001 |
| Genotype x Sex x Dose | F(3,113)=0.52, p=.67 |
| Genotype x Sex x Time Bin | F(11,1243)=0.62, p=.71 |
| Genotype x Dose x Time Bin | F(33,1243)=1.45, p=.09 |
| Sex x Dose x Time Bin | F(33,1243)=1.05, p=.40 |
| Genotype x Sex x Dose x Time Bin | F(33,1243)=1.59, p=.06 |
| *Gad2:Ift88 KO* |  |
| Locomotor activity |  |
| Genotype | F(1,85)=0.72, p=.79 |
| Sex | F(1,85)=3.12, p=.08 |
| Dose | F(3,85)=82.88, p<.001 |
| Time Bin | F(11,935)=56.81, p<.001 |
| Genotype x Sex | F(1,85)=1.02, p=.31 |
| Genotype x Dose | F(3,85)=0.51, p=.68 |
| Genotype x Time Bin | F(11,935)=0.51, p=.72 |
| Sex x Dose | F(3,85)=0.27, p=.85 |
| Sex x Time Bin | F(11,935)=0.69, p=.59 |
| Dose x Time Bin | F(33,935)=12.46, p<.001 |
| Genotype x Sex x Dose | F(3,85)=0.30, p=.83 |
| Genotype x Sex x Time Bin | F(11,935)=0.59, p=.67 |
| Genotype x Dose x Time Bin | F(33,935)=0.61, p=.82 |
| Sex x Dose x Time Bin | F(33,935)=1.45, p=.15 |
| Genotype x Sex x Dose x Time Bin | F(33,935)=1.47, p=.14 |
| Repeated Beam Break |  |
| Genotype | F(1,85)=0.002, p=.97 |
| Sex | F(1,85)=4.39, p=.04 |
| Dose | F(3,85)=34.33, p<.001 |
| Time Bin | F(11,935)=16.73, p<.001 |
| Genotype x Sex | F(1,85)=0.27, p=.59 |
| Genotype x Dose | F(3,85)=0.97, p=.41 |
| Genotype x Time Bin | F(11,935)=1.27, p=.27 |
| Sex x Dose | F(3,85)=0.19, p=.91 |
| Sex x Time Bin | F(11,935)=0.87, p=.52 |
| Dose x Time Bin | F(33,935)=10.93, p<.001 |
| Genotype x Sex x Dose | F(3,85)=0.46, p=.71 |
| Genotype x Sex x Time Bin | F(11,935)=0.75, p=.61 |
| Genotype x Dose x Time Bin | F(33,935)=0.96, p=.50 |
| Sex x Dose x Time Bin | F(33,935)=1.64, p=.048 |
| Genotype x Sex x Dose x Time Bin | F(33,935)=0.90, p=.58 |

**Supplemental Table 2:** Statistical analysis of acute cocaine induced locomotor activity and repeated beam breaks for 1hr following administration for Dat:Ift88 and Gad2:Ift88 mice.

| **Repeated cocaine** |  |
| --- | --- |
| *Dat:Ift88 KO* |  |
| Locomotor activity |  |
| Genotype | F(1,113)=6.61, p=.01 |
| Sex | F(1,113)=0.37, p=.54 |
| Dose | F(3,113)=186.10, p<.001 |
| Day | F(4,452)=72.91, p<.001 |
| Genotype x Sex | F(1,113)=0.98, p=.33 |
| Genotype x Dose | F(3,113)=4.96, p=.003 |
| Genotype x Day | F(4,452)=2.29, p=.07 |
| Sex x Dose | F(3,113)=0.42, p=.74 |
| Sex x Day | F(4,452)=3.53, p=.01 |
| Dose x Day | F(12,452)=8.34, p<.001 |
| Genotype x Sex x Dose | F(3,113)=2.19, p=.09 |
| Genotype x Sex x Day | F(4,452)=0.86, p=.47 |
| Genotype x Dose x Day | F(12,452)=2.27, p=.02 |
| Sex x Dose x Day | F(12,452)=2.64, p=.005 |
| Genotype x Sex x Dose x Day | F(12,452)=0.59, p=.82 |
| Stereotypy |  |
| Genotype | F(1,113)=1.06, p=.31 |
| Sex | F(1,113)=2.05, p=.16 |
| Dose | F(3,113)=26.21, p<.001 |
| Day | F(4,452)=51.57, p<.001 |
| Genotype x Sex | F(1,113)=4.28, p=.04 |
| Genotype x Dose | F(3,113)=2.05, p=.11 |
| Genotype x Day | F(4,452)=0.27, p=.82 |
| Sex x Dose | F(3,113)=1.55, p=.21 |
| Sex x Day | F(4,452)=0.68, p=.54 |
| Dose x Day | F(12,452)=6.01, p<.001 |
| Genotype x Sex x Dose | F(3,113)=0.80, p=.50 |
| Genotype x Sex x Day | F(4,452)=0.34, p=.77 |
| Genotype x Dose x Day | F(12,452)=1.22, p=.29 |
| Sex x Dose x Day | F(12,452)=0.80, p=.60 |
| Genotype x Sex x Dose x Day | F(12,452)=1.16, p=.32 |
| *Gad2:Ift88 KO* |  |
| Locomotor activity |  |
| Genotype | F(1,85)=0.17, p=.68 |
| Sex | F(1,85)=7.52, p=.007 |
| Dose | F(3,85)=70.41, p<.001 |
| Day | F(4,340)=40.27, p<.001 |
| Genotype x Sex | F(1,85)=1.42, p=.24 |
| Genotype x Dose | F(3,85)=1.45, p=.24 |
| Genotype x Day | F(4,340)=1.73, p=.17 |
| Sex x Dose | F(3,85)=0.22, p=.88 |
| Sex x Day | F(4,340)=1.71, p=.71 |
| Dose x Day | F(12,340)=4.87, p<.001 |
| Genotype x Sex x Dose | F(3,85)=0.55, p=.65 |
| Genotype x Sex x Day | F(4,340)=0.25, p=.85 |
| Genotype x Dose x Day | F(12,340)=0.67, p=.72 |
| Sex x Dose x Day | F(12,340)=1.11, p=.36 |
| Genotype x Sex x Dose x Day | F(12,340)=0.48, p=.88 |
| Repeated Beam breaks |  |
| Genotype | F(1,85)=0.13, p=.91 |
| Sex | F(1,85)=5.11, p=.03 |
| Dose | F(3,85)=20.68, p<.001 |
| Day | F(4,340)=31.27, p<.001 |
| Genotype x Sex | F(1,85)=0.21, p=.65 |
| Genotype x Dose | F(3,85)=1.83, p=.15 |
| Genotype x Day | F(4,340)=0.30, p=.88 |
| Sex x Dose | F(3,85)=1.21, p=.31 |
| Sex x Day | F(4,340)=0.30, p=.82 |
| Dose x Day | F(12,340)=6.00, p<.001 |
| Genotype x Sex x Dose | F(3,85)=0.53, p=.66 |
| Genotype x Sex x Day | F(4,340)=1.60, p=.19 |
| Genotype x Dose x Day | F(12,340)=0.71, p=.69 |
| Sex x Dose x Day | F(12,340)=1.00, p=.44 |
| Genotype x Sex x Dose x Day | F(12,340)=0.48, p=.88 |

**Supplemental Table 3:** Statistical analysis of repeated cocaine administration induced locomotor and repeated beam break plasticity following administration for Dat:Ift88 and Gad2:Ift88 mice.

| **Conditioned Place Preference** |  |
| --- | --- |
| *Dat:Ift88 KO* |  |
| Baseline Preference |  |
| Genotype | F(1,11)=0.00, p=.99 |
| Sex | F(1,11)=0.84, p=.38 |
| Genotype x Sex | F(1,11)=0.66, p=.43 |
| Conditioned Place Preference |  |
| Genotype | F(1,11)=8.39, p=.02 |
| Sex | F(1,11)=3.46, p=.09 |
| Genotype x Sex | F(1,11)=0.49, p=.50 |
| Reinstatement |  |
| Genotype | F(1,11)=6.00, p=.03 |
| Sex | F(1,11)=0.26, p=.62 |
| Genotype x Sex | F(1,11)=1.11, p=.31 |
| Extinction |  |
| Genotype | F(1,11)=0.54, p=.48 |
| Sex | F(1,11)=0.35, p=.57 |
| Day | F(9,99)=0.57, p=.66 |
| Genotype x Sex | F(1,11)=0.17, p=.69 |
| Genotype x Day | F(9,99)=1.52, p=.22 |
| Sex x Day | F(9,99)=0.54, p=.68 |
| Genotype x Sex x Day | F(9,99)=0.55, p=.67 |
| *Gad2:Ift88 KO* |  |
| Baseline Preference |  |
| Genotype | F(1,25)=0.53, p=.47 |
| Sex | F(1.25)=5.30, p=.03 |
| Genotype x Sex | F(1,25)=3.10, p=.09 |
| Conditioned Place Preference |  |
| Genotype | F(1,25)=2.41, p=.13 |
| Sex | F(1,25)=9.99, p<.001 |
| Genotype x Sex | F(1,25)=1.41, p=.25 |
| Reinstatement |  |
| Genotype | F(1,25)=0.12, p=.73 |
| Sex | F(1,25)=0.39, p=.54 |
| Genotype x Sex | F(1,25)=1.02, p=.32 |
| Extinction |  |
| Genotype | F(1,25)=4.22, p=.05 |
| Sex | F(1,25)=0.54, p=.47 |
| Day | F(9,99)=1.31, p=.27 |
| Genotype x Sex | F(1,25)=1.78, p=.20 |
| Genotype x Day | F(9,99)=1.61, p=.18 |
| Sex x Day | F(9,99)=1.21, p=.31 |
| Genotype x Sex x Day | F(9,99)=0.86, p=.50 |

**Supplemental Table 4:** Statistical analysis of repeated cocaine administration induced locomotor and repeated beam break plasticity following administration for Dat:Ift88 and Gad2:Ift88 mice.

**Supplemental figures:**

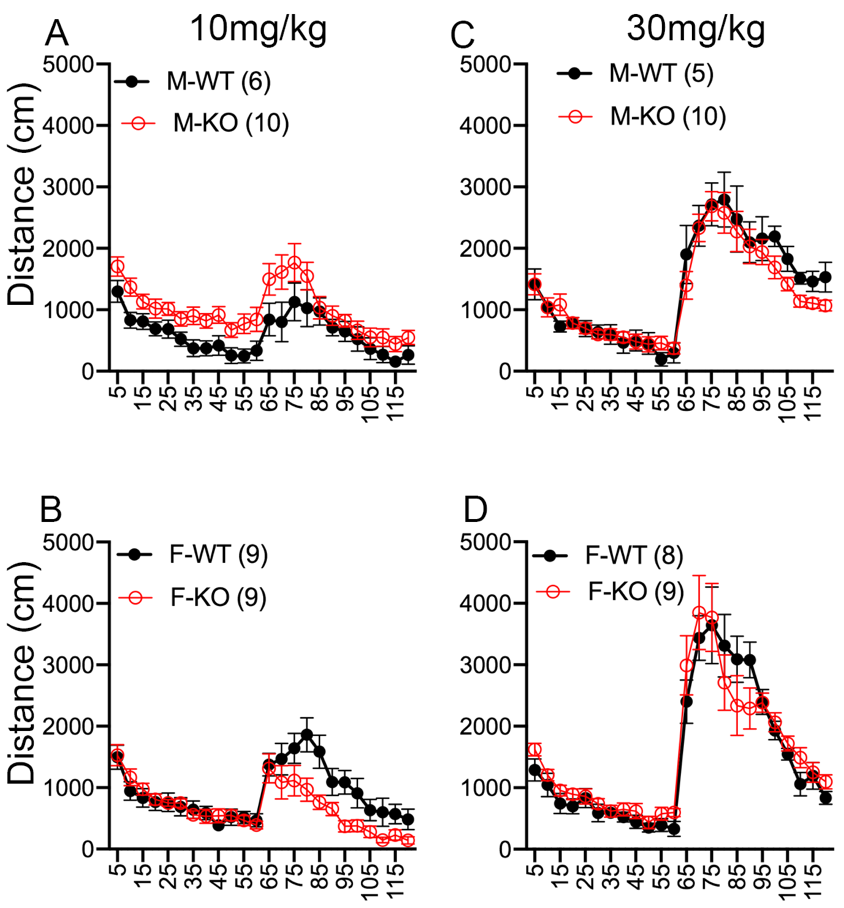

**Supplemental figure 1: Sex differences in acute locomotor responses to cocaine of Dat:Ift88^KO^ mice.** (A-B) Time course of horizontal distance traveled per 5 min bin in male (A) & female (B) WT & KO mice in response to 10mg/kg. There were no main effects of genotype, but there was a genotype x sex x dose interaction (F(3,113)=3.62, p=.02), such that at the 10 mg/kg dose, there was a genotype x sex interaction (F(1,30)=7.24, p=.01). (C-D) Time course of horizontal distance traveled per 5 min bin in male (C) & female (D) WT & KO mice in response to 30mg/kg.

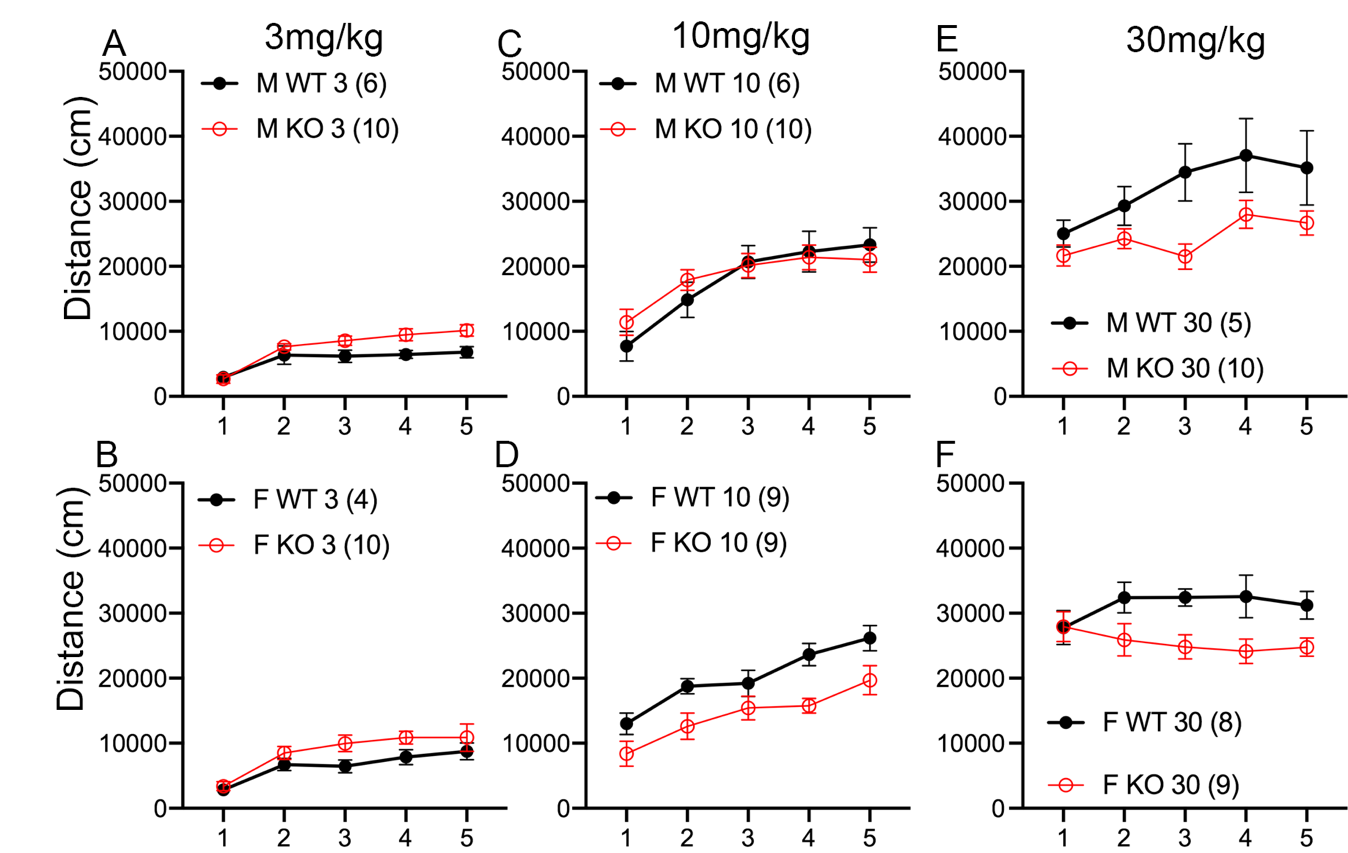

**Supplemental figure 2: Sex differences in locomotor sensitization to cocaine of Dat:Ift88^KO^ mice.** (A-B) Time course of horizontal distance traveled per 5 min bin in male (A) & female (B) WT & KO mice in response to 3mg/kg. There were no main effects of genotype, but there was a genotype x sex x dose interaction (F(3,113)=3.62, p=.02), such that at the 10 mg/kg dose, there was a genotype x sex interaction (F(1,30)=7.24, p=.01). (C-D) Time course of horizontal distance traveled per 5 min bin in male (C) & female (D) WT & KO mice in response to 10mg/kg. (E-F) Time course of horizontal distance traveled per 5 min bin in male (E) & female (F) WT & KO mice in response to 30mg/kg.

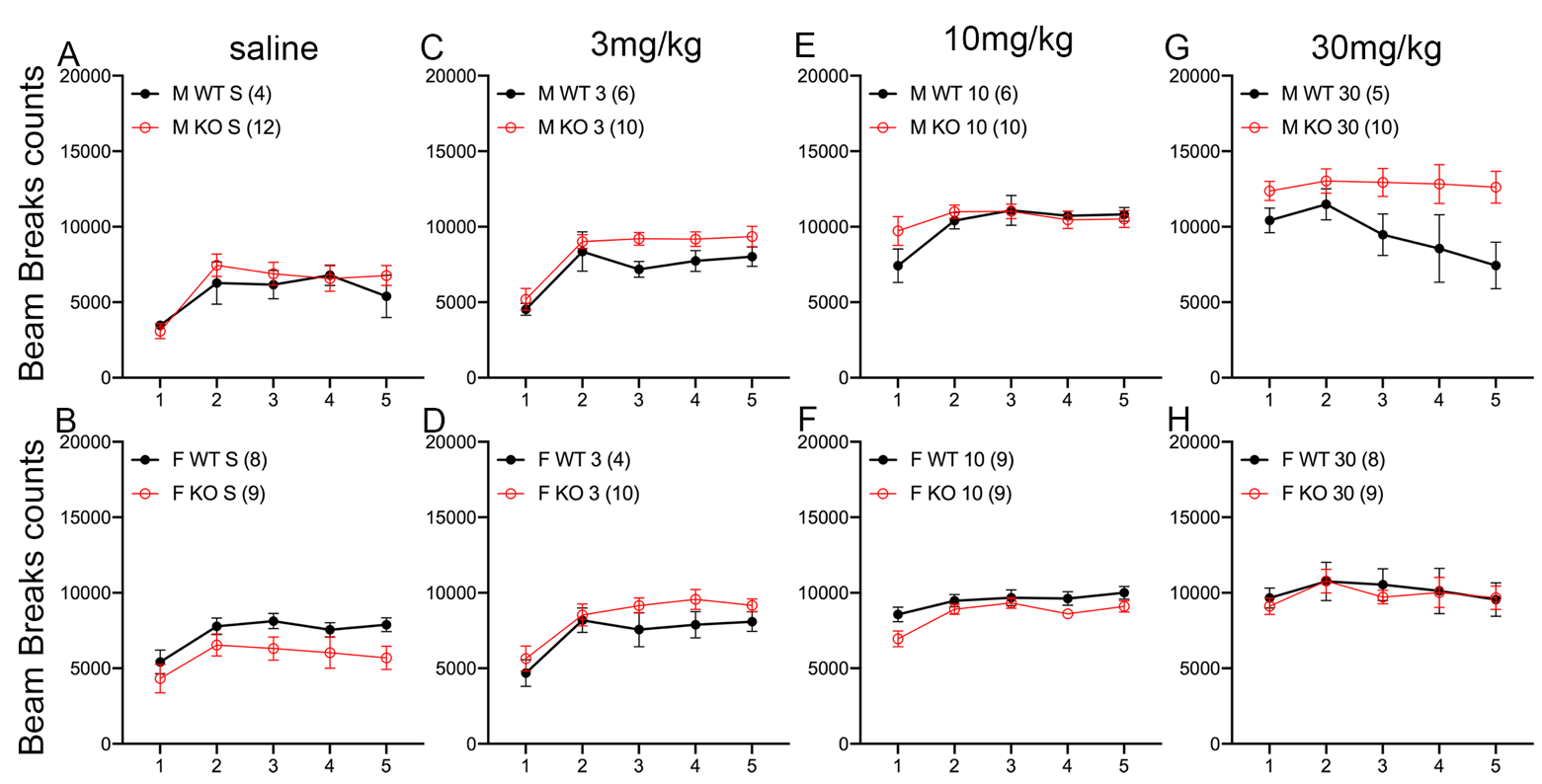

**Supplemental figure 3: Sex differences in repeated beams breaks to cocaine of Dat:Ift88^KO^ mice.** (A-B) Time course of horizontal distance traveled per 5 min bin in male (A) & female (B) WT & KO mice in response to saline. There were no main effects of genotype, but there was a genotype x sex x dose interaction (F(3,113)=3.62, p=.02), such that at the 10 mg/kg dose, there was a genotype x sex interaction (F(1,30)=7.24, p=.01). (C-D) Time course of horizontal distance traveled per 5 min bin in male (C) & female (D) WT & KO mice in response to 10mg/kg. (E-F) Time course of horizontal distance traveled per 5 min bin in male (E) & female (F) WT & KO mice in response to 30mg/kg. (G-H) Time course of horizontal distance traveled per 5 min bin in male (E) & female (F) WT & KO mice in response to 30mg/kg.
